# Disentangling heritability and plasticity effects on *Populus fremontii* leaf reflectance across a temperature gradient

**DOI:** 10.1101/2024.10.21.619129

**Authors:** M. M. Seeley, E. Thomson, G. J Allan, B. C. Wiebe, T. G. Whitham, K. R. Hultine, H. F. Cooper, G. P. Asner, C.E. Doughty

## Abstract

Globally, vegetation biodiversity is expected to decline as the rate of plant adaptation struggles to keep pace with rising temperatures. To support conservation efforts through remote sensing, we disentangled the nested effects of genetic and environmental influences on reflectance spectra, leveraging spectroscopy to assess plant adaptations to temperature. Specifically, we quantified the relative effect of plasticity and heritability on *Populus fremontii* (Fremont cottonwood) leaf reflectance using clonal replicates propagated from 16 populations and grown across three common gardens spanning a mean annual temperature gradient representing the thermal range of *P. fremontii*. We used variance partitioning to decompose phenotypic variation expressed in the leaf spectra into genotypic and environmental components to estimate broad-sense heritability. Heritability was strongly expressed in the spectral red edge (∼680-750nm) and shortwave infrared (∼1400-3000nm), though the heritability peak in the red edge was sensitive to extreme temperatures. By comparing distances of group centroids in principal component space, we determined that *P. fremontii* intraspecific spectral variation was shaped by the interaction between common garden site conditions and source population. Support vector machine models indicated pronounced environmental influence on spectral variation, as *P. fremontii* source population and garden location were classified at 71.8% and 92.6% accuracy, respectively. These findings emphasize the utility of reflectance data in separating genetic and environmental influences on plant phenotypes, offering a pathway to scale these insights across broader landscapes and aid in the conservation and management of vulnerable ecosystems in a warming climate.

## Introduction

Globally, forests are threatened by climate change through increasing temperatures and drought, leading to widespread tree mortalities (Allen et al., 2010). Many regions have already experienced declines in biodiversity, and these declines are predicted to accelerate (Pimm et al., 2014). Remote sensing, specifically imaging spectroscopy, has emerged as a powerful tool for addressing these challenges, offering insights into biodiversity, species distributions, and ecosystem health at the landscape scale (Rose et al., 2015; Seeley & Asner, 2021; Turner et al., 2003). For example, forest conservation applications of imaging spectroscopy include accurate, high-resolution species mapping (e.g. Balzotti et al., 2020; Seeley et al., 2023c), large-scale biodiversity estimations to inform and support protected area establishment (Asner et al., 2017; Féret & Asner, 2014), and forest physiological and chemical monitoring (Brodrick & Asner, 2017; Roberts et al., 2003; Weingarten et al., 2021). Imaging spectroscopy can also be used to explore plastic responses to changing environmental conditions and heritable, intraspecific trait variation that enables species to adapt to changing climates and novel conditions (Czyż et al., 2020; Seeley & Asner, 2023; Wang et al., 2022).

Previous research has successfully linked leaf and canopy spectra to both environmental and genetic factors. For example, visible through shortwave-infrared (VSWIR) reflectance data have been used to track environmental responses, such as changes in leaf traits (e.g. leaf mass per area, chlorophyll) due to stress or climatic variation (Czyż et al., 2020; Seeley & Asner, 2023). It has also been used to distinguish individuals within a species based on their genotype (Blonder et al., 2020; Cavender-Bares et al., 2016; Seeley et al., 2023b). However, many of these studies focus on either genetic or environmental influences, with limited attention to how environmental sensitivity of plant traits, and therefore spectra, can affect phylogenetic studies or how genetic differences in plasticity influence spectral responses to environmental adaptation. Together, these considerations have the potential to reveal insights into how plants adapt and respond to changing conditions.

We used a common garden experiment to assess plastic and genotypically constrained responses of leaf reflectance spectra in *Populus fremontii* (Fremont cottonwood). *P. fremontii* inhabits riparian zones in the southwest United States and Mexico and is recognized as one of the most important foundation species in these regions (Ellison et al., 2005; Whitham et al., 2006) due to its ability to structure communities across multiple trophic levels, drive ecosystem processes, and influence biodiversity via genotypically based functional trait variation (Ferrier et al., 2012; Grady et al., 2013). Despite active planting projects, the areal extent of *P. fremontii* has declined dramatically over the last century due to water extraction, competition with invasive species, and climate change (Braatne et al., 1996; Shafroth et al., 1995; Stromberg, 1998). To support management decisions for *P. fremontii* and protect riparian ecosystems vulnerable to climate change, we attempted to link leaf-level spectral data to temperature adaptations. By deconstructing VSWIR spectra into heritable and plastic components, we aimed to map and monitor *P. fremontii* responses and tolerance to changing climate conditions and to derive a method for quantifying the heritable resilience (intraspecific variation in plant traits that could result in greater ecosystem resilience; Turner, 2023; Vahsen et al., 2023) of *P. fremontii* populations to extreme temperatures at large spatial scales.

Heritability is a statistic that estimates the degree of variation in a phenotypic trait that is due to genetic variation between individuals of a population. Measuring heritability in plant traits has long been a practice in selective breeding programs searching for heritable traits to maximize yield and quality (Bradshaw, 2017). Broad-sense heritability is defined as H^2^=v(G)/v(E), where v(G) is the proportion of genetic variance and v(E) is the proportion of environmental variation, typically quantified using clonal replicates (Hühn, 1975). Thus, broad-sense heritability can vary according to the amount of environmental variation experienced by individuals. As spectroscopy integrates a vast array of leaf traits in its analyses and is a better predictor of plant and ecosystem function than a subset of foliar traits (Yan et al., 2021), quantifying heritability across the reflectance spectrum may yield novel insights into genetic versus environmental controls on leaf spectra, which are linked to functional traits, and reveal parts of the spectrum useful for retrieving phylogenetic information.

By examining *P. fremontii* leaf reflectance across three common gardens spanning the thermal range of *P. fremontii*, we sought to disentangle the genetic and environmental factors shaping its adaptive capacity to varying temperatures. Specifically, our objectives were to: 1) quantify the relative contributions of genetics (source population) and environmental influences (common garden site) to intraspecific spectral variation, 2) explore adaptations to environmental extremes by analyzing reflectance spectra, with a focus on phenotypic plasticity and genetic differentiation under extreme temperatures, and 3) identify conserved and variable regions across the VSWIR spectra of *P. fremontii* across the common gardens. Where we do not specifically assess genetic variation in spectral plasticity, we simplified our general discussion of plasticity here as the degree to which leaf spectra change across different environments, acknowledging that plasticity has a genetic component (Cooper et al., 2019; Laitinen & Nikoloski, 2019; Scheiner, 1993).

## Materials and Methods

### Study Sites

Three common garden sites were established across the thermal range of *P. fremontii* in 2014. Two were located at the thermal edges of the *P. fremontii* temperature range (Yuma, Arizona: 22.8°C and Canyonlands, Utah: 10.7°C; Figure S1), and one garden was in the middle of this range (Agua Fria, Arizona: 17.2°C; Figure 1; Table 1). As Yuma experiences extreme temperatures where ambient temperatures could exceed the critical temperature at which Photosystem II is disrupted (t_crit_) (Moran et al., 2023), we were able to assess the responses of leaf spectra when trees experienced air temperatures approaching and surpassing their critical leaf temperature thresholds (Hultine et al., 2020; Moran et al., 2023). Given the large temperature gradients across the three common garden sites, temperature was expected to be the primary factor influencing physiological processes (Figure 2; Cooper et al., 2019, 2022).

**Figure 1:**
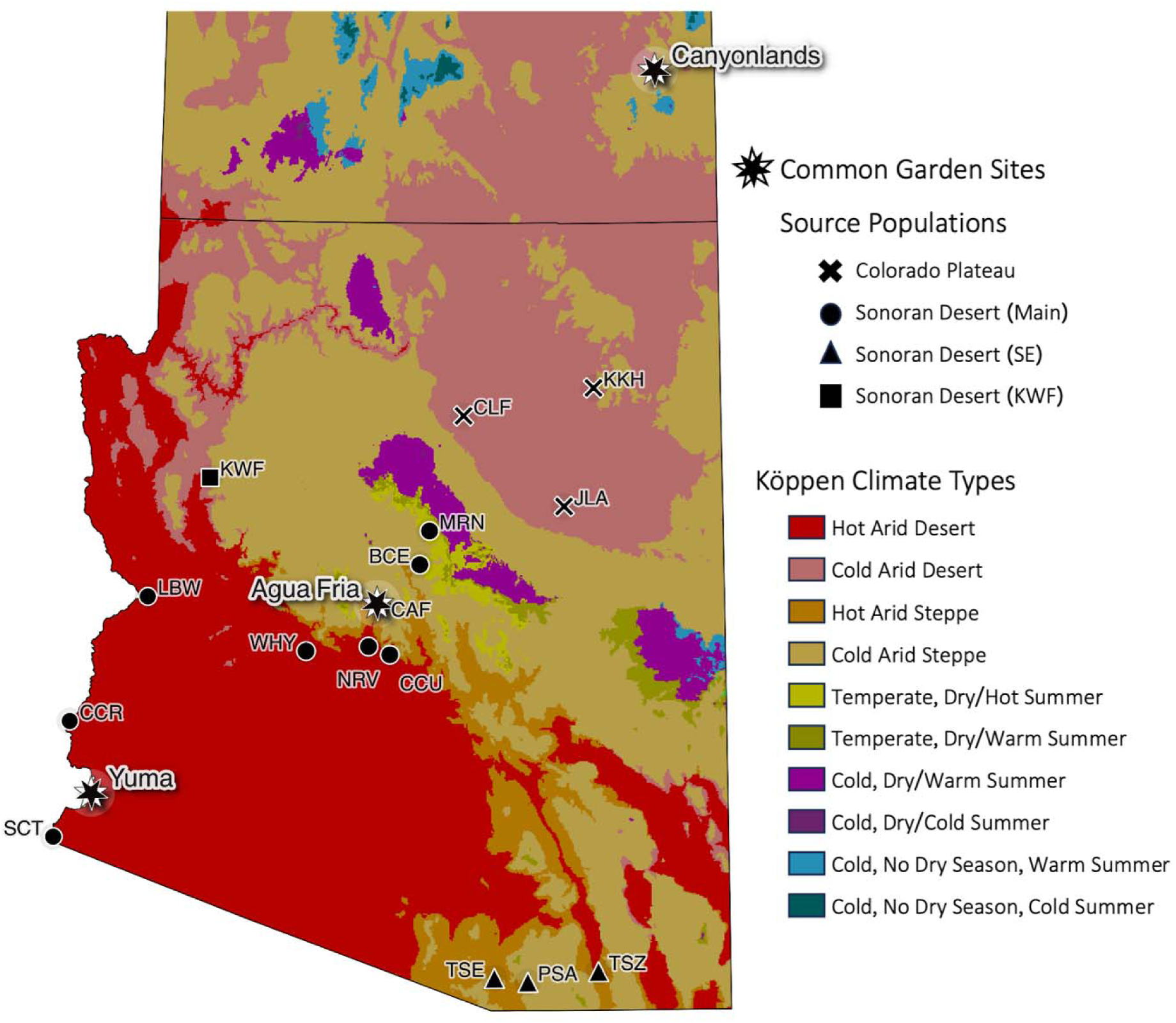
Source population and common garden site locations in Arizona and southern Utah. Background colors represent Köppen climate types.

**Figure 2:**
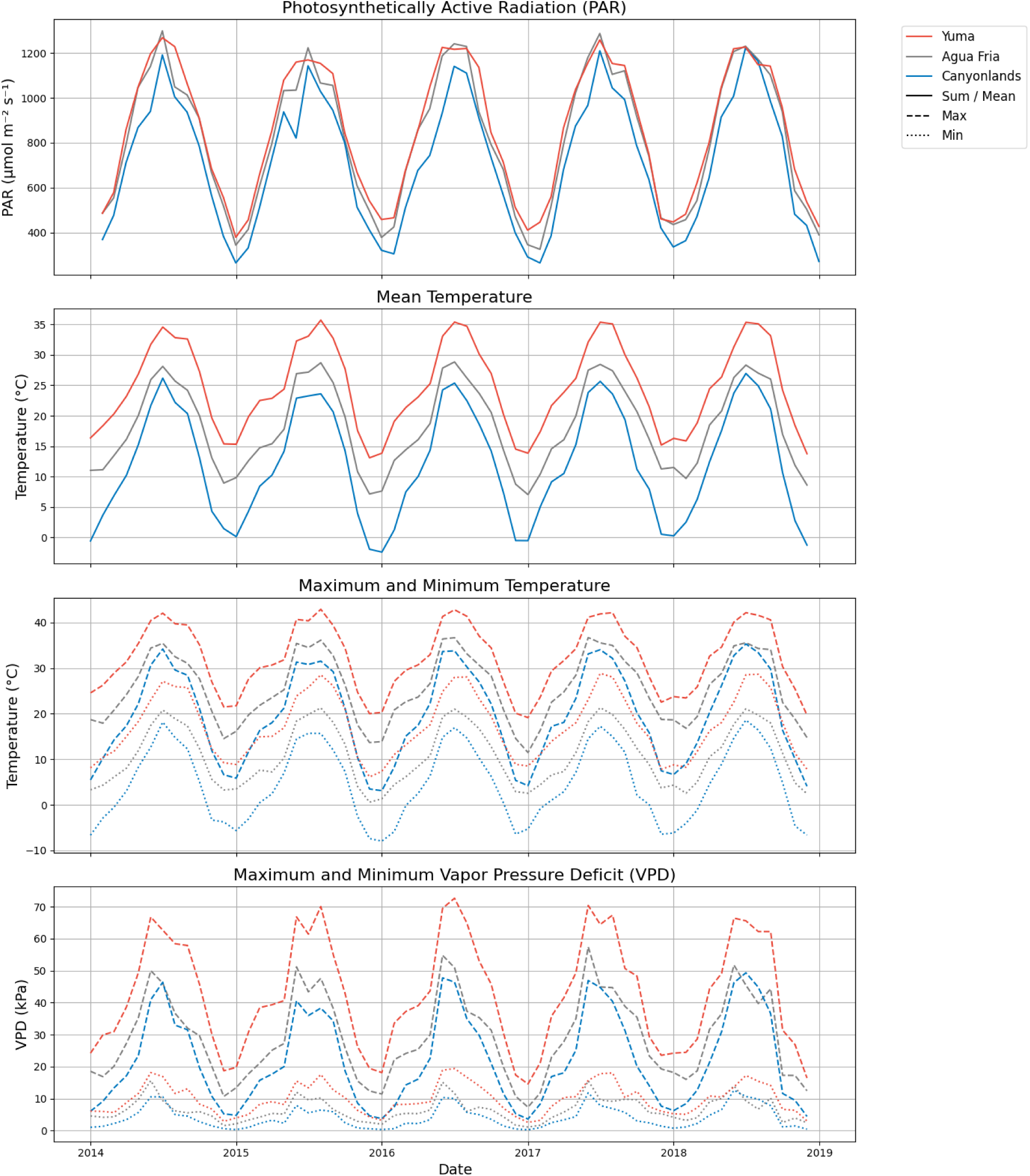
Environmental conditions at the common garden sites from 2014 (garden establishment) to 2018 (data collection). All data represent the mean monthly values. Photosynthetically active radiation (PAR) data are the daily sum radiation values collected every three hours by the Moderate Resolution Imaging Spectroradiometer (MODIS). Mean, minimum, and maximum temperature and vapor pressure deficit (VPD) were obtained from the PRISM Gridded Climate Data.

**Table 1:** Source population and study site environmental conditions. For source populations, the ecotype is listed.

| **Source**<br>**Population** | **Elevation** | **Mean Annual**<br>**Temperature (°C)** | **Mean Annual**<br>**Precipitation (mm)** | **Ecotype** |
| --- | --- | --- | --- | --- |
| MRN | 1774 | 10.4 | 593 | Sonoran Desert Main |
| KKH | 1920 | 10.7 | 258 | Colorado Plateau |
| JLA | 1507 | 12.3 | 212 | Colorado Plateau |
| CLF | 1299 | 14.1 | 176 | Colorado Plateau |
| KWF | 1126 | 15 | 243 | Sonoran Desert KWF |
| BCE | 1109 | 15.6 | 432 | Sonoran Desert Main |
| PSA | 1234 | 15.7 | 471 | Sonoran Desert SE |
| TSZ | 1219 | 16.9 | 322 | Sonoran Desert SE |
| CAF | 988 | 17.2 | 440 | Sonoran Desert Main |
| TSE | 986 | 17.5 | 402 | Sonoran Desert SE |
| WHY | 575 | 19.6 | 284 | Sonoran Desert Main |
| CCU | 696 | 19.9 | 349 | Sonoran Desert Main |
| NRV | 666 | 19.9 | 337 | Sonoran Desert Main |
| SCT | 26 | 22.1 | 88 | Sonoran Desert Main |
| LBW | 143 | 22.3 | 137 | Sonoran Desert Main |
| CCR | 70 | 22.6 | 97 | Sonoran Desert Main |
| **Common Garden Site** |  |  |  |  |
| Canyonlands | 1581 | 10.7 | 225 |  |
| Agua Fria | 998 | 17.2 | 453 |  |
| Yuma | 49 | 22.8 | 93 |  |

Each garden hosted cuttings from sixteen geographically separate (minimum 13.5 km) *P. fremontii* populations across Arizona spanning 12.2°C Mean Annual Temperature. From each cutting, clonal replicates were propagated in a greenhouse. 32 clonal replicates of each cutting were planted in each common garden, across 4 stratified random blocks to minimize the effect of within-garden variation on a population. Each common garden hosted 16 wild populations of *P. fremontii*, eight genotypes (cuttings) from each population (each located >20 m away), and 32 clonal replicates from each genotype, resulting in 4096 *P. fremontii* saplings per garden. Trees at the common gardens were routinely watered to facilitate establishment in the arid southwest. A detailed description of the common garden establishment can be found in Cooper et al. (2019, 2022).

We note that, while all environmental differences between common garden sites are included in the variance explained by site, we assume that the primary factor affecting individuals at each common garden site is temperature following Cooper et al. (2019, 2022).

*P. fremontii* in the common gardens represented two distinct ecotypes, the Colorado Plateau and Sonoran Desert ecotypes (Blasini et al., 2021; Bothwell et al., 2023; Ikeda et al., 2017). *P. fremontii* were collected from sixteen distinct source populations (Table 1), three of which were located on the Mogollon Rim and are part of the Colorado Plateau ecotype (also referred to as the Utah High Plateau/Mogollon Rim ecotype; Blasini et al., 2021; Ikeda et al., 2017). The remaining thirteen source populations were part of the Sonoran Desert ecotype. Within the Sonoran Desert ecotype, three sub-ecotypes (northern and southern) were represented (Bothwell et al., 2023). The larger, northern sub-ecotype includes nine source populations and is referred to as the main Sonoran Desert cluster, while the southeast (SE) Sonoran Desert sub-ecotype includes three populations in the southeastern portion of Arizona. Lastly, one population (KWF) did not fall into either Sonoran Desert sub-ecotype.

### Leaf Measurements

We measured leaf reflectance spectra from trees at the three common gardens in May 2018, four years after the saplings had been planted. Due to the differential survivorship of the saplings, the number of trees sampled from each population at each garden varied (0-52; Table S1). Three mature leaves were collected from each tree, and, where possible, leaves were harvested from different branches at the same leaf position on the shoot, preferentially just below the new growth, to standardize for development. Only fully expanded, sunlit adult leaves were chosen. Leaves were harvested by hand and transported in a cool, dark box to the nearby field station for processing the same day.

Spectral measurements were collected using an Analytical Spectra Devices (ASD) leaf clip, plant probe, and field spectrometer from 350 to 2500 nm at 1-nm intervals (Analytical Spectra Devices Inc., Boulder, CO, USA). Three measurements were taken on the adaxial surface of each leaf in the center and averaged. The spectrometer was optimized using a Spectralon white reference taken after every three leaves. To minimize noise, wavelengths with a low signal-to-noise ratio were removed (<400 nm and >2450 nm). Brightness normalization was applied to all spectral measurements to minimize variation not related to leaf chemistry and structure (Kruse et al., 1993; Myneni et al., 1989).

### Leaf trait measurements and analyses

Following spectral measurements, leaves were scanned using a digital scanner (Canon LiDE 110) to determine their area, oven dried at 70 ◦C for three days and weighed for their dry mass. Leaf mass per area (LMA) was calculated by dividing the leaf’s dry mass (g) by its fresh one-sided area (m^2^).

We next calculated solar-weighted reflectance, or albedo, for the visible (400-700nm) and near-infrared (NIR)-shortwave infrared (SWIR; 700-2500nm). Reflectance spectra (not brightness normalized) were weighted by the solar irradiance spectrum (ASTM G173-03 Reference Spectra derived from SMARTS version 2.9; Gueymard, 2004) to calculate broadband albedo (Bartlett et al., 2011) according to Equation 2, where λ represents each wavelength:

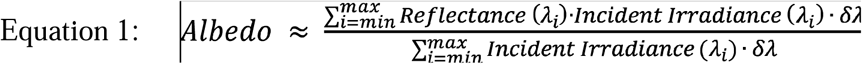

Lastly, we estimated chlorophyll content using the chlorophyll index – green (CIG) according to Equation 2.

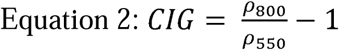

*ρ*_800_ is the reflectance at 800□nm (in the NIR) and *ρ*_500_ is that at 550□nm (green). As chlorophyll absorbs green light, higher CIG values indicate greater leaf chlorophyll content. To determine whether the common garden sites induced changes in leaf traits, a Tukey-Kramer multiple comparison test from the scipy python package (Virtanen et al., 2020) was applied to the albedo, LMA, and CIG.

### Phenotypic variation in leaf spectra

To explore phenotypic variation in leaf spectra, we first visualized spectral variation across the three common gardens. Mean source population spectra at each common garden site were plotted and colored according to site. Spectral differences between the same populations at different common garden sites were further visualized using principal component analysis (PCA). PCA reduces the dimensionality of the spectral data from over 2000 dimensions to a handful of dimensions that explain the primary spectral variation across multiple channels (Boardman & Green, 2000; Green & Boardman, 2000; Thompson et al., 2017). We randomly selected 104 spectra from among the five populations that were represented in all three common gardens (CLF, KWF, BCE, PSA, and SCT). Using scikit-learn (Pedregosa et al., 2011), we applied PCA to spectra data, plotting the first two principal components (PC) along the x and y axes. As higher PC are often important in spectroscopy applications for discriminating differences in surface chemistry and other traits (Dai et al., 2022; Thompson et al., 2017), we further visualized spectral differences across ecotypes and common gardens in the first 20 PCs. As the effect of extreme heat at Yuma was obvious in the first two PCs (Figure 4), this second analysis included spectral data from only Agua Fria and Canyonlands. Data were grouped by ecotype (Colorado Plateau, Sonoran Desert Main, and Sonoran Desert Southeast (SE)) and garden, randomly selecting spectra from 40 trees from each ecotype-garden combination. We then calculated the centroids (group mean) of each ecotype-garden combination in the first 20 PCs and measured the Euclidean distances between these centroids. Distances were then projected into 2D space using multidimensional scaling (MDS), following the approach of Seeley et al. (2023c).

### Decomposing genotypic and environmental variation across the spectra and estimating broad-sense heritability

To quantify the relative effects of environmental versus genotypic variation across the VSWIR, we used variance partitioning to decompose observed variance at each wavelength into the relative contribution of site, ecotype, population, genotype, or clone. All sampled data was used across all common garden sites. Following the method of Asner et al. (2014) and Fyllas et al. (2009), variance partitioning was calculated using a linear mixed-effects model for each wavelength across the VSWIR using the lme4 *R* package (Bates et al., 2015). The model was fitted using Residual Maximum Likelihood (REML). All effects were treated as random. Common garden site was included as a crossed random effect, while clones were nested within genotypes, which were nested within populations, which were nested within ecotype. The total spectral variance about the mean for a given wavelength was, therefore, quantitatively parsed into the variance explained by site (σ2_s_), ecotype (σ2_e_), populations within ecotype (σ2_p_), genotypes within populations (σ2_g_), clones within genotypes (σ2_c_) and residual variance (σ2_r_) according to Equation 3:

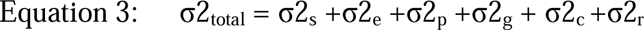

We next calculated broad-sense heritability across the VSWIR. Heritability at each wavelength was calculated using the classic broad sense heritability equation (H^2^ = V_G_/V_P_), where V_G_ is genotypic variation, and V_P_ is total variation (V_G +_ V_E_). Variance explained by ecotypes, populations, and genotypes is categorized as genotypic variance (V_G_), as the genetic material within these groups differs. Variance explained by site and clones, as well as residual variance, is categorized as environmental variation (V_E_), or spectral plasticity, as the genetic material within these groups is identical, so any observed differences in the spectra must be due to environment or measurement error. Broad-sense heritability was plotted across the VSWIR spectrum. This analysis was applied first to data from all three sites and then to data from Canyonlands and Agua Fria alone. To obtain confidence intervals for H^2^, we employed a nonparametric bootstrap approach. Bootstrapping, a method commonly used for obtaining uncertainty estimates for heritability (Schweiger et al., 2016), involves resampling the dataset with replacement and recalculating the variance with each subsampled dataset. We performed 1000 iterations for each wavelength, sampling 31 spectra from each common garden x ecotype combination, resulting in n=279 for each iteration. Mean H^2^ of the 1000 iterations were presented along with the 95% confidence interval calculated from the bootstrapped estimates.

### Using leaf spectra to predict genotypic and environmental information

To explore whether leaf spectra could be used to extract both genotypic and environmental information, we used a support vector machine (SVM) to predict *P. fremontii* source population (genotypic variation) and garden location (environmental variation). SVMs efficiently handle highly dimensional datasets by creating decision boundaries in n-dimensional space that maximize the distance between class boundaries and sample points (Melgani & Bruzzone, 2004). SVMs tend to outperform other machine learning algorithms such as random forest for species and genotype classifications when applied to spectroscopy data (e.g. Dalponte et al., 2012). For each radial basis function kernel SVM, we optimized hyperparameter selection through a cross-validation grid search implemented in python with the scikit-learn package (Pedregosa et al., 2011). As with the PCA analyses, 104 samples from each garden were randomly selected from the same five source populations (CLF, KWF, BCE, PSA, and SCT). For all models, data were divided into training/test datasets using a 70/30 stratified random split.

To assess the generalizability of models in unseen and extreme environments in predicting genotypic information, we trained three additional SVM models. Each model was trained to predict the source population using *P. fremontii* spectra from two of the gardens (n = 208) and tested on unseen spectra from the third garden (n = 104). We repeated this experiment three times, with a different garden withheld each time. This allowed us to compare the applicability of the model in extreme environments and assess how well the model can extrapolate outside of its training data.

## Results

VSWIR leaf spectra responded to common garden treatment, particularly at Yuma, which represents the upper end of the cottonwood thermal range. Brightness-normalized reflectance of all populations at the Yuma site diverged from that at both Canyonlands and Agua Fria (Figures 2a & 3a). Albedo in both the visible and NIR-SWIR differed across the common gardens (p<0.05) and was highest at Yuma. In the visible, albedo was lowest at Canyonlands, and in the NIR-SWIR, albedo was lowest at Agua Fria. CIG, indicative of chlorophyll content, was highest in Canyonlands (mean = 4.8) and Agua Fria (mean = 4.7). Yuma had the lowest CIG, with a mean of 2.8 (p<0.05).

LMA, an important driver of spectral variation, was highest in the Yuma common garden and lowest in Canyonlands (p<0.05; Figure 4b). Mean LMA across all populations at Canyonlands was 72.9 g/m^2^ with a standard deviation of 10.2 g/m^2^. At Agua Fria, the mean LMA was 88.1 g/m^2^, though it was the most variable with a standard deviation of 19.1 g/m^2^. Yuma exhibited the highest LMA at 108.4 g/m^2^ and a standard deviation of 11.2 g/m^2^. In addition to differing across the gardens, LMA also varied among source populations; a two-way ANOVA revealed significant main effects of both garden environment and population on LMA (p<0.05 for both factors).

In PC space, spectra differed across the common gardens and among the ecotypes (Figure 4). The first two PC of spectra from all common gardens explained 57.8 and 35.3% of the variance, respectively. These components primarily separated Yuma spectra from those at Canyonlands and Agua Fria (Figure 4a). When comparing spectra from different ecotypes at Canyonlands and Agua Fria, the Colorado Plateau ecotype separated from both Sonoran Desert ecotypes regardless of common garden in the first 20 PCs. Within the Sonoran Desert ecotypes (Main and SE), common garden was the primary driver of spectral divergence in PC space.

### Variance partitioning and broad-sense heritability

Across all gardens, spectral variance was highest in the visible (∼400-680 nm) and red-edge wavelengths (∼680-750 nm). When Yuma was included in the analysis, variance in these wavelengths was driven largely by the common garden site (Figures 2 and 3), and heritability in these regions was correspondingly low with a maximum heritability of 0.08 for the visible (Figure 5). The red-edge heritability peak present when Yuma was excluded collapsed with the inclusion of Yuma, leaving a smaller heritability peak at the longer wavelengths. Additionally, site-level variation in the visible (∼400-680 nm) and red-edge wavelengths (∼680-750 nm) was minimal without Yuma. Instead, variance at these wavelengths was driven more by genetic factors (population and genotype), with corresponding high spectral heritability (∼0.36 and ∼0.66, respectively).

**Figure 3:**
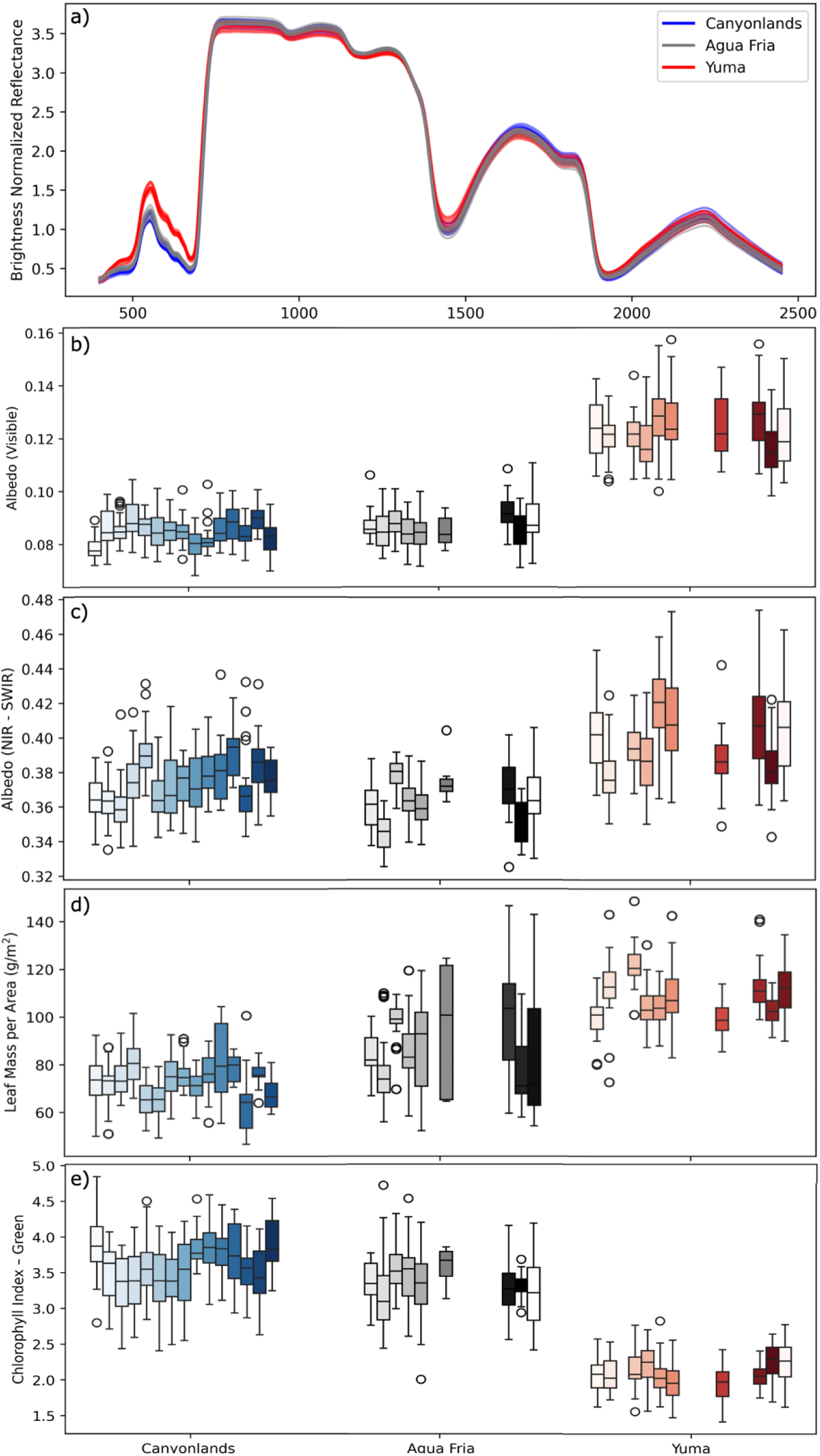
a) Mean brightness normalized reflectance of each source population, colored by common garden site. See Figure S2 for reflectance prior to brightness normalization. Boxplots of albedo calculated across the b) visible and c) near-infrared-shortwave infrared, d) leaf mass per area (LMA), and e) chlorophyll index – green (CIG). Within each garden, boxplots are ordered according to source population mean annual temperature, with cooler populations on the left and warmer populations on the right. Gaps correspond to populations that were not sampled at a given garden. Boxplots represent a different number of sampled trees per population per site.

**Figure 4:**
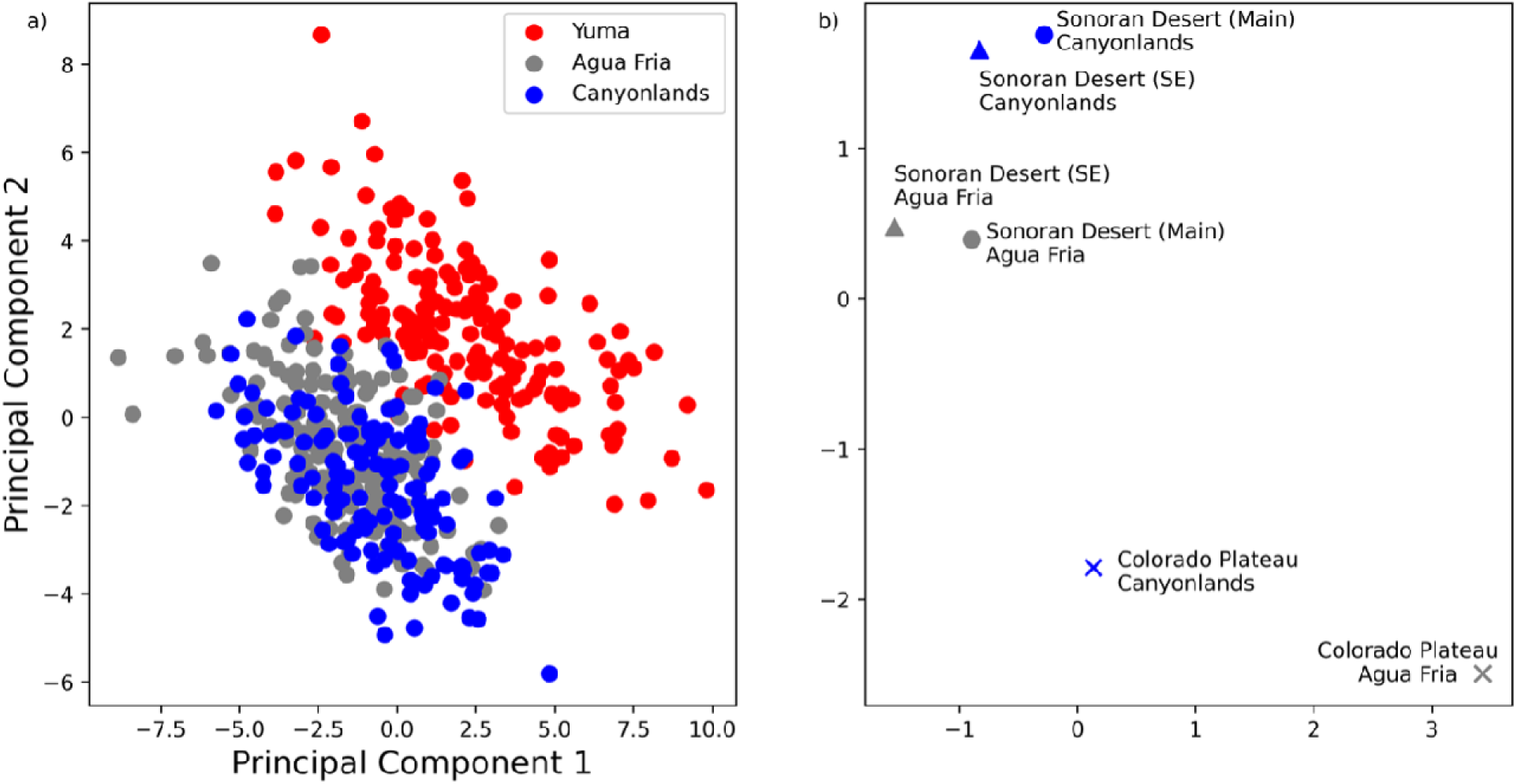
a) First and second principal components (PC) of leaf spectra colored according to the common garden site. b) Distance between the centroids of source population grouped by ecotype x common garden in PC space. Colors represent common garden (blue: Canyonlands, grey: Agua Fria). Shapes correspond to the ecotype (x: Colorado Plateau, circle: Sonoran Desert, triangle: Sonoran Desert SE).

**Figure 5:**
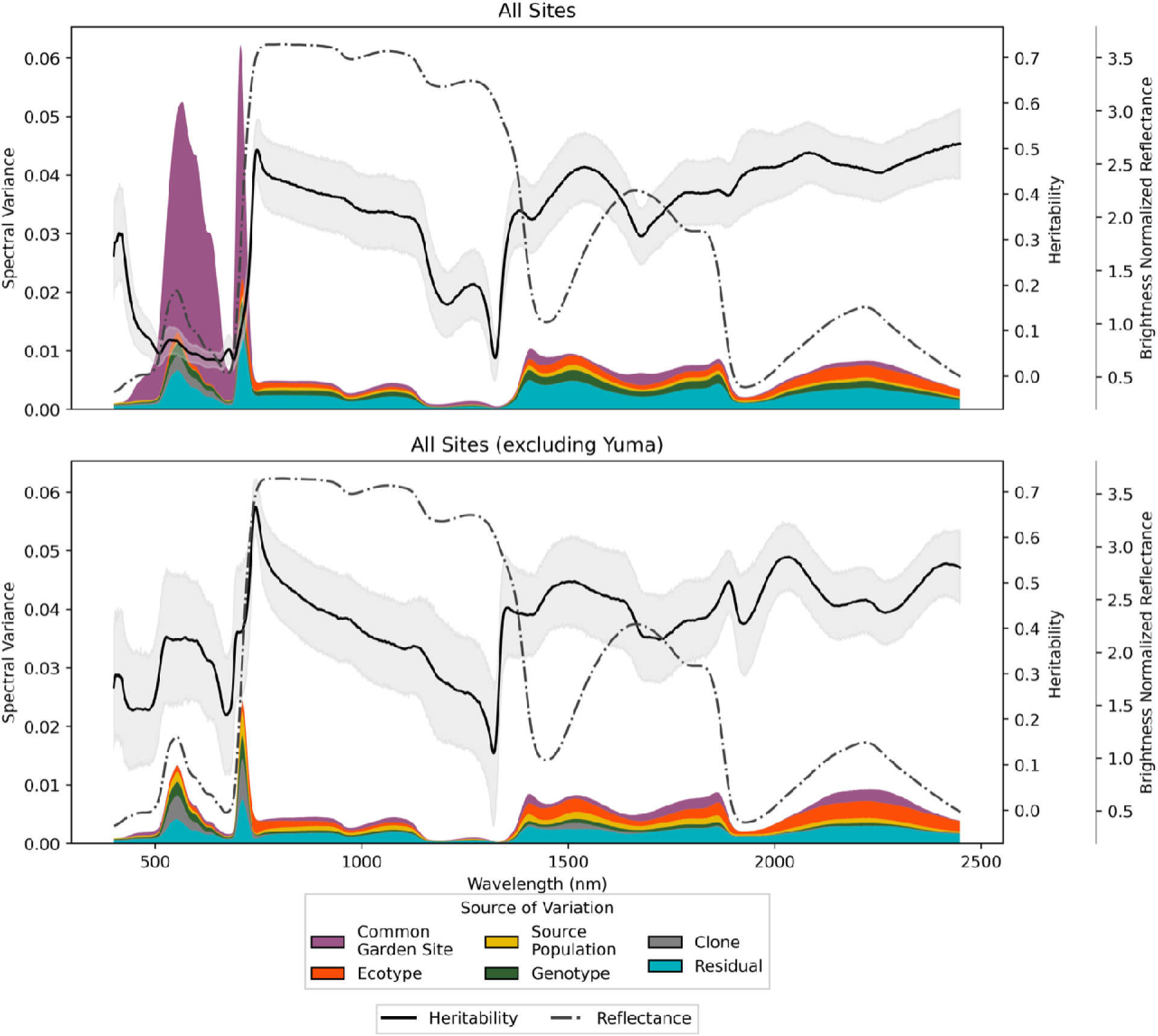
Heritability of reflectance spectra (solid line) and its 95% bootstrapped confidence interval (grey fill), mean brightness normalized reflectance spectra (dotted line), and the source of variability (filled colors) from the visible to shortwave infrared for *P. fremontii* at a) all sites and b) Canyonlands and Agua Fria.

Across all gardens, heritability in *P. fremontii* reflectance was greatest in the SWIR (1400 – 2450 nm) and the transition from red-edge to NIR (∼750 nm), regardless of the inclusion of Yuma (Figure 5). In the shorter wavelengths of the NIR (∼750-1140 nm) and SWIR, site-level variation explained less of the variance than genotypic factors in general, and the spectral differences caused by Yuma were relatively minimal, resulting in high heritability (max: 0.51; min: 0.31). When Yuma was excluded, heritability peaked in the red-edge (740 nm; heritability: 0.67). Heritability had a negative relationship with wavelength in the NIR until ∼1320 nm where heritability increased and remained high in the SWIR, although it varied across this region (max: ∼0.56, min: ∼0.37). Across the VSWIR, particularly heritable regions of the spectra included the red-edge/NIR transition (∼750 nm) and the SWIR. More environmentally sensitive (plastic) regions included the visible and red-edge as well as much of the NIR.

### SVM analysis and leaf predictions

SVM models were able to discriminate between spectra based on both common garden site and source population at 92.6% and 71.8% accuracy, respectively (Figure 6). The SVM trained to classify spectra based on the common garden site (environmental variation) outperformed the SVM trained to discriminate spectra based on source population (genotypic variation). The common garden site SVM accurately predicted spectra from the Yuma site 100% of the time; confusion in this model occurred primarily between Canyonlands and Agua Fria, with the true positive rate for Canyonlands and Agua Fria being 94% and 84%, respectively. For the source population SVM, the true positive rate ranged from 61% for the KWF population to 84% for CLF. When predicting source population using one common garden site as a holdout for testing, model performance declined (Figure S4). Source populations at Canyonlands were predicted with 38.5% accuracy when trained on data from Agua Fria and Yuma. At Agua Fria and Yuma, accuracy was 37.5% and 31.7%, respectively, when SVM models were trained on data from the other common garden sites.

**Figure 6:**
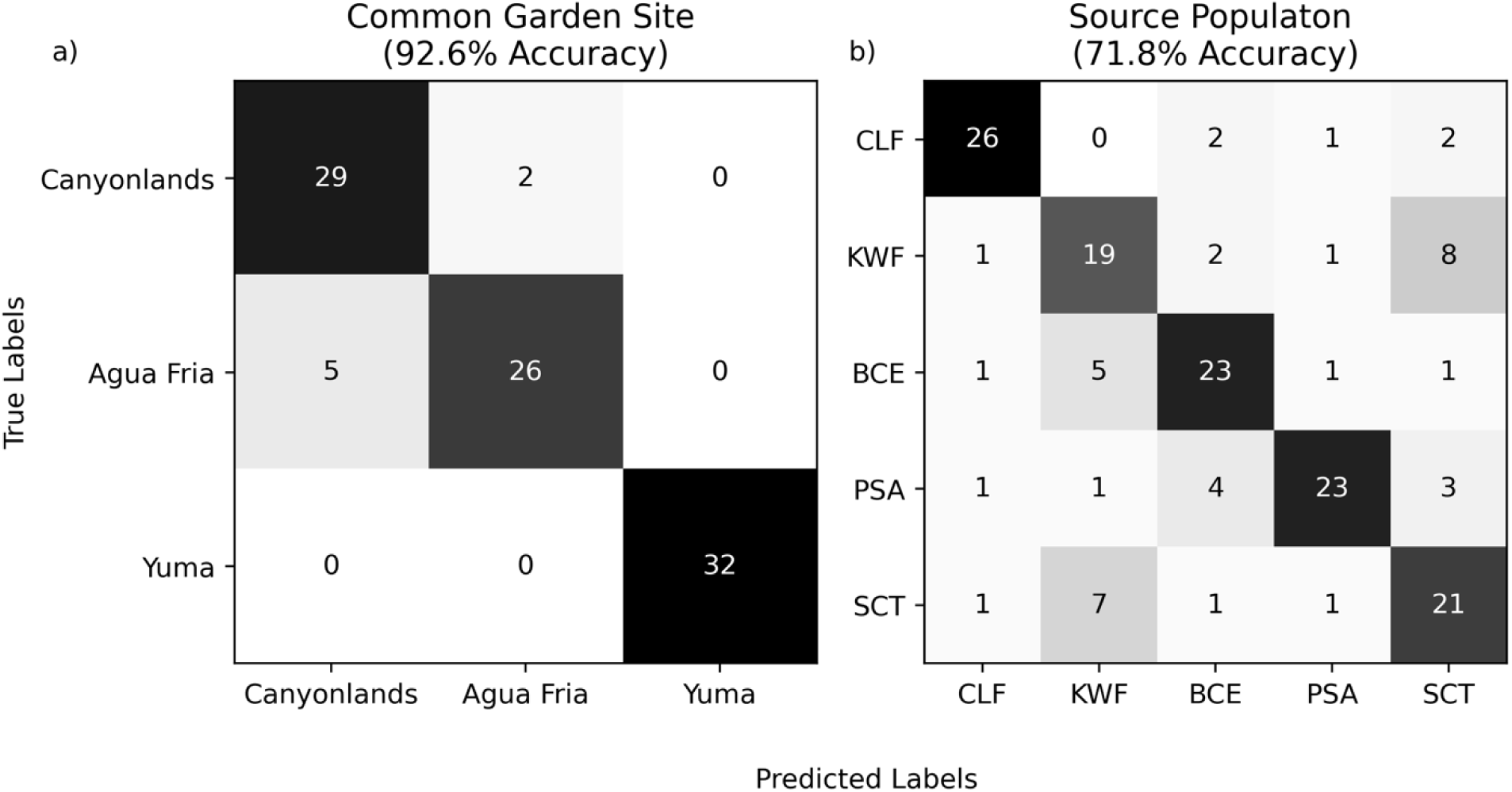
Support vector machine confusion matrix results for predicting a) common garden site and b) source population. Source populations are ordered left to right and top to bottom from lowest to highest mean annual temperature.

## Discussion

High temperatures (>50°C) induced physiological stress in *P. fremontii,* altering their spectral signatures. The environmental variability introduced by Yuma, the hottest common garden, resulted in lower heritability in the visible and red edge regions of the reflectance spectra, discussed further below. We confirmed that trees growing at Yuma experienced thermal stress, as indicated by the environmental data and lower CIG. Moran et al. (2023) documented temporal changes in leaf thermal tolerance (t_crit_) at Yuma, confirming that *P. fremontii* undergoes physiological adjustments in response to exposure to extreme temperatures. Further, Sankey et al. (2020) demonstrated GxE interactions among *P. fremontii* at the Agua Fria common garden, observing differences in canopy temperature between genotypes using thermal imagery from unmanned aerial vehicles. Plastic responses at the individual level can include increased LMA (observed at Yuma), increased heat shock proteins (HSP) to help repair heat damage, concentrations of which have been found to correlate with LMA (Knight & Ackerly, 2003), trichome development to promote self-shading (Banowetz et al., 2008), an accumulation of phenolics and isoprenoids for their ability to protect photosynthetic apparatuses and antioxidant properties (Bartwal et al., 2013; Smirnoff, 2005), and increased production of soluble sugars (Wahid & Close, 2007) and saturated fatty acids (Ahmad & Prasad, 2011; Tang et al., 2016). In addition to the physiological changes, Cooper et al. (2019) identified phenological (e.g. bud set, bud flush) differences between clones located at different gardens; for example, bud flush occurred earlier at the Yuma garden.

As many of the physical and chemical changes such as increased LMA, plant phenols, soluble sugars, and fatty acids can be estimated using spectroscopy (Akkaya, 2018; Asner & Martin, 2009, 2016), it is unsurprising that *P. fremontii* at Yuma differed spectrally from those at the other sites. For *Panax ginseng*, differences between VSWIR reflectance of heat-stressed and control plants corresponded to spectral regions linked to fatty acids, sugars, protein, chlorophyll, and nitrogen (Park et al., 2021). Similarly, other studies have found differences between control and heat-stressed plants using fluorescence (Faqeerzada et al., 2023) and visible-NIR (Kim et al., 2022) spectroscopy. While plastic responses to heat stress may decrease spectral heritability, environmentally sensitive regions of the spectra offer opportunities to understand physiological responses to extreme temperatures at large scales and may serve as environmental indicators of stress events. Yuma is near the southern edge of the distribution of *P. fremontii* where it may not survive future climatic conditions (e.g., Blasini et al., 2021; Moran et al., 2023), so identifying environmental indicators of extreme environmental conditions may allow us to monitor stress at the shifting range edge of *P. fremontii*.

### Visible and Red Edge (400-750nm)

The visible and red-edge regions demonstrated the greatest sensitivity to extreme temperatures. Without Yuma, heritability estimates in the visible red-edge regions were higher, suggesting that genetic factors played a stronger role in explaining trait variation under non-stress conditions. Heritability in the visible and red edge has been documented across many species (Čepl et al., 2018; Czyż et al., 2020; Feng et al., 2017; McManus et al., 2016). For example, *Metrosideros polymorpha* varieties that organize along environmental gradients diverged in chlorophyll content, which was associated with differences in the visible wavelengths (Seeley et al. 2023b). Feng et al. (2017) found that the red edge was predictive of rice genotypes, and both Čepal et al. (2018) and Czyż et al. (2020) observed a red edge signal in the heritability of *Pinus sylvestris* (Scotts pine) and *Fagus sylvatica* (European beech) leaf reflectance, respectively. In heat-stressed conditions, however, increased physiological plasticity, such as changes in chlorophyll concentration, can reduce the expression of genetic differences (Posch et al., 2024; Seeley et al., 2025b; Stone et al., 2018), thereby lowering heritability.

The inclusion of Yuma, characterized by elevated temperatures, likely introduced greater environmental variance and potentially GxE interactions, altering the proportion of phenotypic variance attributable to genetics. Although we did not directly measure chlorophyll concentration, both the visible and red-edge regions are closely associated with chlorophyll and other photosynthetic pigments (Horler et al., 1983; Vogelmann et al., 1993). Further, the CIG indicated that chlorophyll content at Yuma was lower than that at the other gardens. Heat stress can lead to disruption of the photosystems, characterized by chlorophyll degradation and changes in pigment ratios (Ivanov et al., 2017; Mathur et al., 2014; Tafesse et al., 2022), which can reduce absorption in the visible range and shift the red-edge position (Tirado et al., 2021; Xie et al., 2019). Reflectance of trees at Yuma were consistent with these processes, as they exhibited higher visible reflectance, similar to that reported for heat-stressed rice (Xie et al., 2019) and maize (Tirado et al., 2021). The change in reflectance suggests physiological changes that both serve as a mechanism of photosynthetic apparatus protection and as are a side effect of the stress response.

### Near-Infrared (750-1400nm)

Heritability across the NIR declined with increasing wavelength in *P. fremontii* and was slightly lower with the inclusion of Yuma. NIR reflectance is primarily influenced by the internal structure of leaves, particularly the spongy mesophyll (Gausman, 1974). LMA, which influences NIR reflectance (Asner et al., 2011; Gara et al., 2021; Serbin et al., 2019), increased across the common gardens and varied among source populations, indicating that both genetic and environmental factors affected LMA. Knight and Ackerly (2003) similarly found elevated LMA under thermal stress in desert plants, linking it to enhanced photosynthetic recovery and identifying high LMA as a convergent trait in plants adapted to hot, arid environments, like *P. fremontii*. The increase in LMA in response to thermal stress at Yuma may have contributed to the decline in heritability in the NIR, though this effect was less pronounced than in the visible region. Heritability in the NIR remained relatively high, however, perhaps because other traits that could further reduce heritability, such as leaf water content (Gausman, 1974; Seeley et al., 2025b), were controlled across the gardens. The heritability peaks we observed in the NIR are consistent with those reported in *Pinus sylvestris* and *Fagus sylvatica* (Čepl et al., 2018; Czyż et al., 2020), though *P. sylvestris* showed an additional region of high heritability between 1130–1330 nm that corresponded to low heritability in *P. fremontii*. Given that much of the NIR signal reflects internal leaf structure (Knipling, 1970; Slaton et al., 2001), these interspecific differences may arise from contrasting leaf morphologies—broadleaf vs. needleleaf.

### Shortwave-infrared (1400-2500nm)

In the SWIR, heritability remained relatively high regardless of the inclusion of the hotter Yuma garden, suggesting that genetic control over traits expressed in the SWIR was robust to environmental variation. The SWIR region is associated with structural and biochemical leaf components such as lignin, phenols, and cellulose (Asner & Martin, 2016), which are influenced by both species-level differences and intra-specific genetic variation (Asner & Martin, 2009; Seeley et al., 2023). These traits are often under stronger genetic regulation and/or less plastic in response to temperature stress compared to traits captured in other spectral regions (Novaes et al., 2010), likely explaining the stability of heritability in this SWIR. Previous studies have likewise identified wavelengths in the SWIR as having high phylogenetic signal (Czyż et al., 2020; McManus et al. 2016; Meireles et al. 2020).

While reflectance in the SWIR has a strong genetic component, this region is also important for separating heat-stressed and control plants, as noted by Park et al. (2021). This is because temperature-dependent leaf traits (e.g. ribulose bisphosphate carboxylation rate) affect reflectance in the SWIR (Park et al., 2021; Serbin et al., 2012), as was demonstrated by Serbin et al. (2012) in two *Populus* species, *P. tremuloides* and *P. deltoides* (Serbin et al., 2012). As LMA is also expressed in the SWIR (Asner et al., 2011; Gara et al., 2021; Serbin et al., 2019), this trait may further contribute to the sensitivity of the SWIR to heat stress. Further, SWIR-associated traits can span a broad heritability range; for example, some phenolic compounds exhibit strong genetic determination, whereas others show considerable plasticity in response to environmental stressors such as heat (Klaper et al., 2001; Rivero et al., 2001). In *P. fremontii,* site-level variation between Agua Fria and Canyonlands was primarily observed in the SWIR, emphasizing the importance of using the full spectrum to distinguish between temperature treatments.

### Albedo

Across the entire VSWIR spectra, albedo increased in response to extreme temperature, likely lowering canopy temperatures and helping mitigate further heat stress. In the NIR – SWIR, albedo decreased from Canyonlands to Agua Fria, supporting the results from Doughty et al. (2018), where albedo declined across a temperature gradient in Peru. Regardless of which region of the VSWIR was analyzed, albedo at Yuma was higher than at the other common gardens. While additional evidence is required to confirm whether the increase in leaf albedo is a universal plant response to extremely high temperatures, this phenomenon aligns with plant ecophysiological principles; many stress factors, including heat, as was demonstrated here, are known to decrease chlorophyll concentration (Carter & Knapp, 2001).

### Heritability and plasticity of reflectance spectra

Intraspecific variation in *P. fremontii* reflectance spectra was driven by GxE interactions, with extreme temperatures at the Yuma site playing a dominant role. Ambient temperatures at Yuma, which approach the upper thermal tolerance limit of *P. fremontii* (Moran et al., 2023), likely caused a threshold change in spectral expression and heritability, as evidenced by the heritability estimates, discussed above, and PCA analyses. In the two PCA scenarios, with and without Yuma, spectra separated according to garden and ecotype, respectively, demonstrating hierarchical effects of environmental and genetic factors. Similar findings were reported by Li et al. (2023), who observed that PC1 of *Nicotiana attenuata* spectra was primarily driven by common garden location, while genotype structured variation along PCs 2–4. Here, the extreme temperatures at Yuma induced leaf physiological responses (e.g. changes in chlorophyll content, LMA) that dominated the spectral signal. This dominant effect of environmental on spectra was further supported by our classification models: SVMs trained to predict garden site achieved ∼14% higher accuracy than those trained to distinguish source populations. Cavender-Bares et al. (2016) similarly found that water treatment was classified more accurately than population origin for *Quercus oleoides*.

When we trained SVMs on data from two common garden sites and tested on the third, classification accuracy for predicting *P. fremontii* source population declined by over 34%, despite controlling for genetic identity. This decline aligns with previous studies demonstrating that the environmental sensitivity of spectra limits large-scale species classifications (e.g., Seeley et al., 2024; 2025a). While *in situ*, ecological mechanisms behind species classification limitations may include distance decay (Rocchini, 2007), intraspecific spectral variation (Czyż et al., 2020; Seeley & Asner, 2023; 2025a), and interspecific trait convergence (Rodríguez-Alarcón et al., 2022), this common garden framework allowed us to control for genetic identity and isolate environmental effects on classification accuracy. The sharp drop in model performance under novel garden conditions suggested that plasticity in spectral traits, arising from site-level environmental variation, was a key limiting factor in model transferability.

Below the physiological temperature threshold at Yuma, spectral differences were predominantly governed by genetic factors, with environmental conditions playing a significant role in shaping phenotypic expression within genetically similar groups. Excluding Yuma in the PCA resulted in the source population (genotype) being the primary driver of separability, observed by the separation of the Sonoran Desert and Colorado Plateau ecotypes in PC space. However, within finer sub-ecotype categories, differences were primarily driven by the common garden, reinforcing the interconnected roles of genetics and environment in shaping spectral traits.

## Conclusions

Reflectance spectra of *P. fremontii* contain valuable information about both the environmental conditions in which they grow and their genetic identities. Intraspecific spectral variation of *P. fremontii* was shaped by both common garden site and source population, with environmental conditions playing a dominant role under extreme stress, such as the high temperatures at the Yuma site. At the upper thermal threshold of *P. fremontii*, the spectral response was most pronounced in the visible and red-edge regions, corresponding to documented heat stress responses. In these regions, spectral heritability decreased when Yuma was considered in the analysis, suggesting that extreme environmental conditions may drive a threshold shift in spectral expression. However, genetic information was retained, and heritability was more consistent in parts of the NIR and SWIR.

These results underscore the potential for using VSWIR reflectance data to disentangle genetic and environmental contributions to plant phenotypes, providing a path forward for scaling these insights to larger landscapes. As airborne and satellite-based imaging spectroscopy data continue to become more widely available, they offer a powerful opportunity to assess phenotypic plasticity and heritable resilience of *P. fremontii* to extreme conditions across broad spatial scales. Spectroscopy may be a powerful tool for monitoring the adaptive capacity of this foundation species in the face of rapid environmental change, supporting efforts to preserve and manage at-risk ecosystems in a warming climate.

## Declarations

### Author contributions

Original idea: CED, ET, GPA; common garden development common gardens: GJA, TGW, KCG, KRH, and HFC; data collection: ET; statistical analysis: MMS, ET, methodology: MMS, ET, CED, BCW; writing the manuscript: MMS, ET. All authors reviewed several drafts and agreed with the final version.

### Availability of data

All data files will be made available on the Figshare database upon acceptance of the manuscript at DOI: 10.6084/m9.figshare.25719585.

### Code availability

Code is available on GitHub and is maintained by Seeley (2025) https://github.com/MegsSeeley/temperature_cottonwood.

