## Supplemental Information for "Disentangling heritability and plasticity effects on *Populus fremontii* leaf reflectance across a temperature gradient"

Table S1: Number of trees sampled at each garden for each source population.

|  | Number of Trees Sampled | | |
| --- | --- | --- | --- |
| Source Population | Canyonlands | Agua Fria | Yuma |
| MRN | 20 |  | 20 |
| KKH | 20 |  | 20 |
| JLA | 35 | 14 |  |
| CLF | 48 | 36 | 20 |
| KWF | 37 | 31 | 47 |
| BCE | 42 | 49 | 41 |
| PSA | 27 | 55 | 40 |
| TSZ | 20 |  |  |
| CAF | 20 | 7 |  |
| TSE | 20 |  |  |
| WHY | 19 |  | 19 |
| CCU | 11 |  |  |
| NRV | 20 |  |  |
| SCT | 16 | 45 | 56 |
| LBW | 19 | 8 | 20 |
| CCR |  | 46 | 51 |

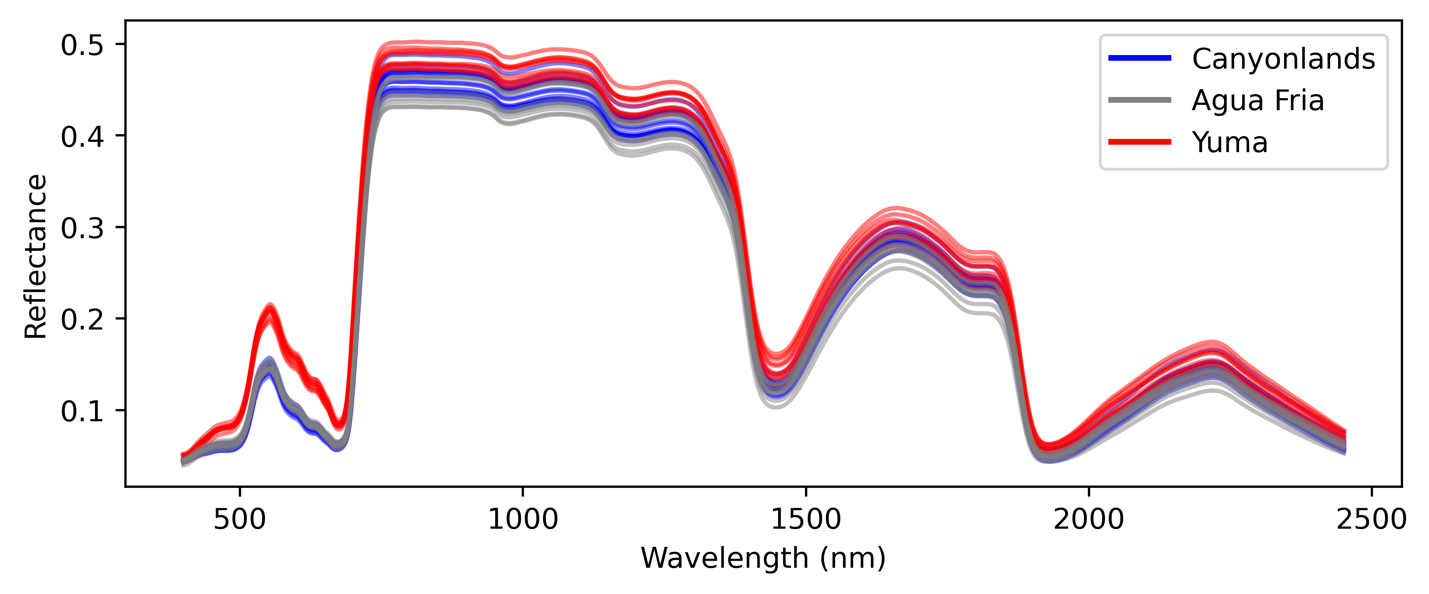

Figure S1: Mean reflectance prior to brightness normalization of each source population at each of the three common garden sites.

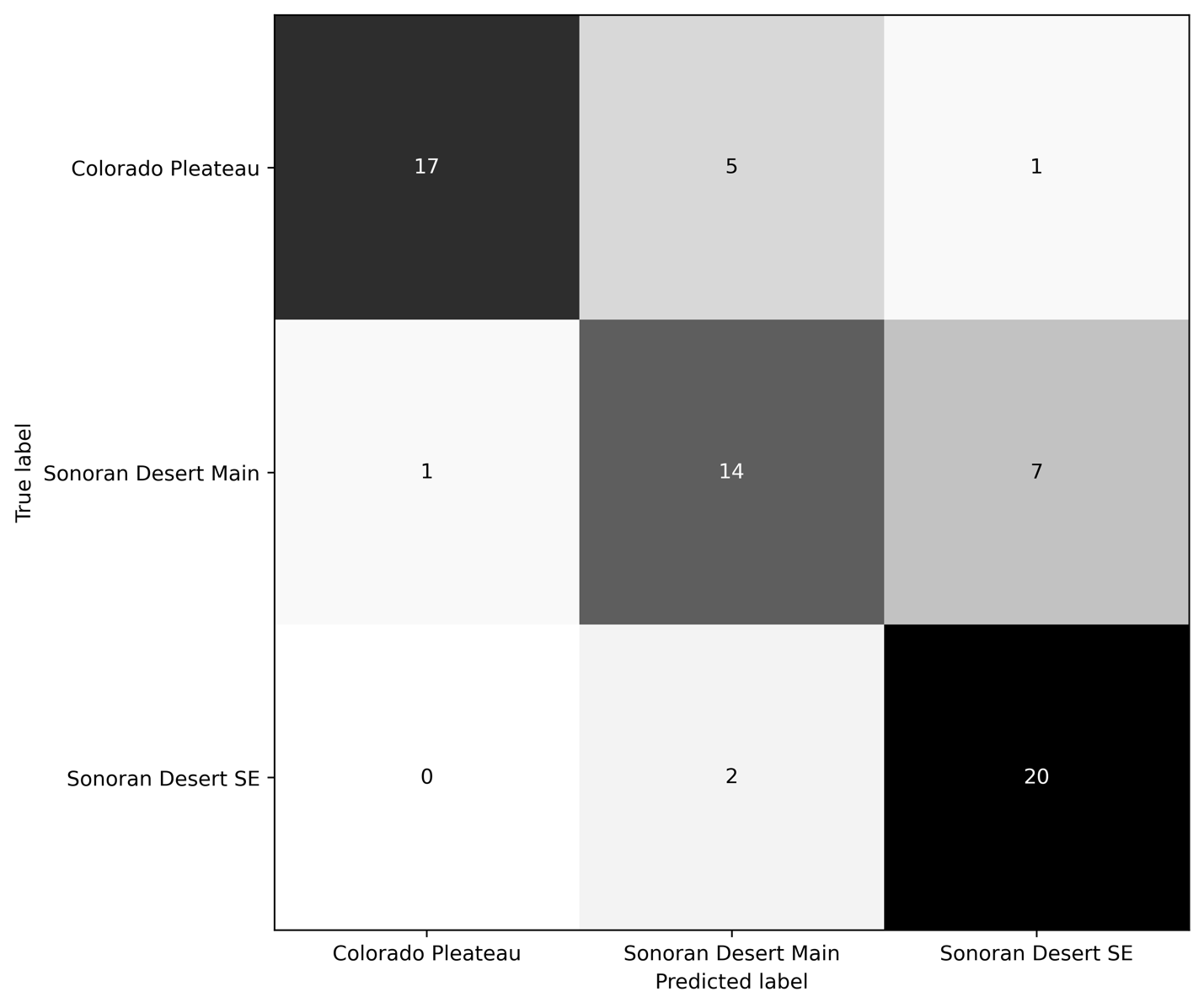

Figure S2: Support vector machine confusion matrix results for predicting ecotype at Agua Fria and Canyonlands.
